# Siliplant1 (Slp1) protein precipitates silica in sorghum silica cells

**DOI:** 10.1101/518332

**Authors:** Santosh Kumar, Nurit Adiram-Filiba, Shula Blum, Javier Arturo Sanchez-Lopez, Oren Tzfadia, Ayelet Omid, Hanne Volpin, Yael Heifetz, Gil Goobes, Rivka Elbaum

## Abstract

- Silicon is absorbed by plant roots as silicic acid. The acid moves with the transpiration stream to the shoot, and mineralizes as silica. In grasses, leaf epidermal cells called silica cells deposit silica in most of their volume by unknown mechanism.
- Using bioinformatics tools, we identified a previously uncharacterized protein in sorghum (*Sorghum bicolor*), which we named Siliplant1 (Slp1). Silica precipitation activity *in vitro*, expression profile, and activity in precipitating biosilica *in vivo* were characterized.
- Slp1 is a basic protein with seven repeat units rich in proline, lysine, and glutamic acid. A short peptide, repeating five times in the protein precipitated silica *in vitro* at a biologically relevant silicic acid concentration. Raman and NMR spectroscopies showed that the peptide attached the silica through lysine amine groups, forming a mineral-peptide open structure. We found Slp1 expression in immature leaf and inflorescence tissues. In the immature leaf active silicification zone, Slp1 was localized to the cytoplasm or near cell boundaries of silica cells. It was packed in vesicles and secreted to the paramural space. Transient overexpression of Slp1 in sorghum resulted in ectopic silica deposition in all leaf epidermal cell types.
- Our results show that Slp1 precipitates silica in sorghum silica cells.

## Introduction

Grasses are well known for their high silica (SiO_2_ ·nH_2_O) content, reaching up to 10% of their dry weight (Hodson *et al*., 2005). Silicon is available to plants as mono-silicic acid [Si(OH)_4_] whose concentration in soil solution usually varies between 0.1 to 0.6 mM (Epstein, 1994). Grass roots actively uptakes silicic acid from soil (Ma *et al*., 2006, 2007) of which more than 90% is loaded into the xylem. A cooperative uptake of silicic acid by two root transporters leads to Si supersaturation in the grass xylem sap (Sakurai et al., 2015). Si concentration may reach 5-7.5 mM, which is 3 to 4 times higher than its saturation concentration (Ma *et al*., 2002; Casey *et al*., 2003; Mitani & Ma, 2005). Silicic acid molecules then move with water inside the plant body to reach the shoots where they are unloaded from the xylem (Yamaji *et al*., 2008). Finally, silicic acid polymerises as solid biogenic silica at several locations inside the plant body. The most prominent sites of silica deposition in grasses are the walls of leaf epidermal cells, abaxial epidermal cells of the inflorescence bracts (glumes and lemma) and root endodermal cells. The silicification mechanism across the cell types is not uniform. Models of deposition suggest either spontaneous formation as a result of evapo-transpirational loss of water, or a tightly controlled process (Kumar *et al*., 2017b). Uptake of silicic acid and its deposition are also affected by plant mechanical damage (McLarnon *et al*., 2017), and possibly other unknown physiological factors affecting root silicon transporters (Talukdar *et al*., 2019).

One of the cell types most frequently silicified in grass leaves is silica cells (Motomura *et al*., 2000; Kumar & Elbaum, 2018). Silica cells are specialized epidermal cells occurring mostly as silica-cork cell pairs, both above and below the leaf longitudinal veins (Kaufman *et al*., 1985), on internode epidermis (Kaufman *et al*., 1969) and on the abaxial epidermis of glumes (Hodson *et al*., 1985). We earlier estimated that the whole process of leaf silica cell silicification is completed within 10 hours of cell division (Kumar & Elbaum, 2018). Within this time period, almost the entire cell lumen is filled with solid silica (Hodson *et al*., 1985). Silica cell silicification is immediately followed by cell death (Kumar & Elbaum, 2018) which ceases the silicification process (Markovich *et al*., 2015; Kumar *et al*., 2017a; Kumar & Elbaum, 2018). Silicification in silica cells is thus different than silicification in macro-hairs and abaxial epidermal cells in lemma where silica deposition takes place in a course of weeks on the thickened cell wall material (Hodson *et al*., 1984; Perry *et al*., 1984), or in glume prickle hairs and papillae where silica is deposited in the empty lumen of these cells even after cell death (Hodson *et al*., 1985). The template on which silica cells deposit silica is unknown. Cell wall polysaccharides such as mixed-linkage glucan (Fry *et al*., 2008; Kido *et al*., 2015) and callose (Law & Exley, 2011; Brugiére & Exley, 2017; Kulich *et al*., 2018) have been suggested as a template for silicification in plants. However, lumen silica structure of *Triticum durum* silica cells has continuously distributed organic matter with the N/C ratio indicative of amino acids (Alexandre *et al*., 2015). This suggests protein(s) as the templating organic matrix for the silicification process in silica cells.

Silicification is widespread among living beings from unicellular microbes to highly evolved multicellular organisms (Perry, 2003). Several bio-silica associated proteins have been reported, for example, silaffins (Kröger *et al*., 1999; Poulsen & Kröger, 2004), silacidins (Wenzl *et al*., 2008) and silicanin-1 (Kotzsch *et al*., 2017) from diatoms; and silicateins (Shimizu *et al*., 1998) and glassin (Shimizu *et al*., 2015) from sponges. Among plants, a short peptide derived from an inducible proline-rich protein precipitates silica *in vitro*. The protein, involved in systemic acquired resistance in cucumber, may precipitate silica locally at attempted fungal penetration sites (Kauss *et al*., 2003). All the above-mentioned protein groups do not share sequence homology.

To study silica deposition in silica cells, we used sorghum (*Sorghum bicolor*), a member of the grass (Poaceae) family. Sorghum is categorized as an active silica accumulator (Hodson *et al*., 2005; Coskun *et al*., 2018). In grasses, leaves appear successively (Skinner & Nelson, 1995). Even within an individual leaf, there is a gradient of cell maturation. The leaf epidermal cell division zone is confined to the base of the leaf and the newly divided cells displace the maturing cells away from the leaf base. Hence, leaf epidermal cells that are close to the leaf apex are older than the cells close to the base (Skinner & Nelson, 1995).

We earlier found that silica cell silicification is confined to elongating leaves, in a well-defined active silicification zone (ASZ) (Kumar *et al*., 2017a; Kumar & Elbaum, 2018). The mineralization initiates in the paramural space of viable silica cells, producing a thick silica cell wall, and restricting the cytoplasmic space to smaller and smaller volumes (Kumar *et al*., 2017a; Kumar & Elbaum, 2018). During this fast process, the cell maintains cell-to-cell connectivity to the neighbouring cells through plasmodesmata (Kumar *et al*., 2017a). As the silicic acid resides in the apoplast in supersaturation, a crucial stage would be the initiation of controlled silica deposition. A possible way would be by adding to the cell wall a biomineralizing protein in appropriate time and place. Here, we report on a previously uncharacterized protein that is expressed in silica cells, exported to the cell wall during their silicification, and induces precipitation of silica *in planta*. Hence, we named this protein as Siliplant1 (Slp1).

## Materials and Methods

### Plant material, growth conditions and tissue nomenclature

Seeds of *Sorghum bicolor* (L.) Moench (line BTx623) were surface sterilized and grown in soil as reported previously (Kumar & Elbaum, 2018). Unless indicated otherwise, we used sorghum seedlings of about 2 weeks of age for our studies. Immature, silicifying leaves (L_IS_) in our present study is analogous to leaf-2, as reported in our earlier studies (Kumar & Elbaum, 2017, 2018; Kumar *et al*., 2017a). The immature leaf was cut into five equal segments. The middle segment is most active in terms of silica cell silicification (Kumar *et al*., 2017a; Kumar & Elbaum, 2018) and was named active silicification zone (ASZ). The segment just older than ASZ (towards leaf tip) was named ASZ+1, while the segment just younger than ASZ (towards leaf base) was ASZ-1. The youngest segment (at leaf base) was named ASZ-2. Mature leaves were cut into 10 equal segments and only the eighth segment from the leaf-base was used for all the experiments.

### Sequence-based analyses of Slp1

Secondary structure prediction was done using the program GOR4 (Combet *et al*., 2000). Intrinsically disordered tertiary structure was predicted using IUPred (Dosztányi *et al*., 2005a,b). The prediction type we chose was ‘long disorder’. Residues with score greater than 0.5 were regarded disordered. SignalP was used to predict signal peptide in Slp1 (Petersen *et al*., 2011). TargetP was used to predict whether the protein is secretory in nature (Emanuelsson *et al*., 2000). The organism group selected was ‘plant’ and the program was run with default cutoff selection.

### Silica precipitation using Peptide-1 and Peptide-3

We tested the peptide sequence HKKPVPPKPKPEPK (Peptide-1) which appears 5 times in Slp1 primary sequence for its silica precipitating activity *in vitro*, and compared it to a mutated peptide, where all lysine groups were replaced by alanine (HAAPVPPAPAPEPA, Peptide-3). We freshly prepared 1 M silicic acid solution by mixing 150 μl of tetramethyl orthosilicate to 850 μl of 1 mM HCl for four minutes under gentle stirring.

For small-scale silica precipitation at 90.9 mM silicic acid, 5 μl of 1 M silicic acid solution was mixed with 50 μl of peptide solution (1.5 or 2.0 mg ml^−1^ in 0.1 M potassium phosphate buffer, pH 7.0) and incubated for 5 minutes under gentle shaking at room temperature. For silica precipitation at 5 mM silicic acid, 55 μl of freshly prepared 1 M silicic acid solution was diluted in 945 μl of 0.1 M potassium phosphate buffer (pH 7.0), of which 10 μl was mixed with 100 μl of Peptide-1 solution (1.5 mg ml^−1^ in 0.1 M potassium phosphate buffer, pH 7.0) and incubated for 30 min under gentle shaking condition. The same reactions lacking any peptide served as control. Sediment was collected by centrifugation at 14000 g for 5 minutes. The supernatant was thrown, and the pellet was washed thrice with 1 ml H_2_O.

For large-scale precipitating of circa 100 mg of silica we incubated 30 mg of peptide (1.5 mg ml^−1^ in 0.1 M potassium phosphate buffer, pH 7.0) with 2 ml of freshly prepared 1 M silicic acid solution for 30 min at room temperature under gentle shaking condition. The precipitate with Peptide-1 was collected by centrifugation at 5000 X g for 5 minutes and washed thrice by H_2_O. The precipitate with Peptide-3 was collected by centrifugation at 5000 X g for 20 min and washed thrice with H_2_O. Both the precipitates were dried at 60 °C.

### Scanning electron microscopy (SEM)

The pellet of the small-scale reaction was suspended in H_2_O and 1 μl of the suspension was smeared on carbon tape, air dried and coated with iridium. The coated sample was observed in Jeol JSM IT100 (Peabody, MA, USA) scanning electron microscope (SEM).

### High resolution transmission electron microscopy (HRTEM)

Powder from peptide-1 large-scale precipitation was resuspended in double distilled water and spread on a TEM grid. HRTEM measurements were carried out on a JEM 2100 JEOL microscope. The samples were imaged at an electron-beam accelerating voltage of 200 kV. The images were collected to characterize the silica internal structure.

### Raman spectroscopic observation of Peptide-1 and silica precipitated with Peptide-1

Lyophilized peptide solution was reconstituted in double distilled water (50 mg ml^−1^) and 1 μl of this solution was put on a steel slide and dried at 37 °C for 30 minutes before measuring the Raman spectra.

The small-scale precipitation of 90.9 mM silicic acid with peptide-1 was dried at 60 °C until attaining a constant dried weight. The dried powder was mounted on a steel slide. Raman spectra of the samples were collected using InVia microspectrometer (Renishaw, New Mills, UK) equipped with polarized 532 nm laser (45 mW max. intensity, 4 μm^2^ beam) excitation under a 50X air objective lens (NA=0.75). Spectra analysis was done in WiRE3 (Renishaw), including smoothing, background subtraction, and peak picking.

### Solid state NMR spectroscopy

Solid state NMR spectra of the peptide-1 large-scale precipitation were recorded on a Bruker 11.7 T Avance III spectrometer (Billerica, MS, USA) equipped with a 4 mm VTN CPMAS probe at a spinning rate of 10 kHz. The ^29^Si cross polarization experiment was done with 8192 scans and recycle delay of 6 sec and the ^29^Si direct excitation experiment was recorded with 2560 scans and recycle delay of 60 sec. Analysis of the ^29^Si spectrum was done using the DMFIT program (Massiot *et al*., 2002). ^13^C cross polarization was done with 2048 scans using recycle delay of 5 sec.

### Tissue-specific expression of Slp1 in sorghum

Polyclonal antibodies against Peptide-1 was raised in rabbit (GenScript, NJ, USA). Crude proteins from root tissues, whole of immature silicifying leaves, mature leaves and immature inflorescence (expected to emerge from the flag leaf of mature plants within about a week) were extracted in the extraction buffer (0.1 M potassium phosphate buffer, pH 7.0; 1% protease inhibitor cocktail, Sigma and 0.1% 2-Mercaptoethanol). Forty-eight μg of the extracted protein from each sample was loaded in separate wells and run on 15% polyacrylamide separating gel under denaturing condition. The gel was blotted onto nitrocellulose membrane and the membrane was put in blocking buffer (5% milk powder in Tris buffer saline, 0.1% Tween 20) for one hour. We incubated the membrane with purified polyclonal antibody against Peptide-1 (1 μg ml^−1^) at 4 °C for overnight. The membrane was washed and incubated with secondary antibody (anti-rabbit IgG mouse monoclonal antibody, Genscirpt, NJ, USA) conjugated with Horseradish peroxidase. Chemiluminescence was developed (PerkinElmer, Akron, US) and the membrane was scanned for one second exposure time.

### Transcript abundance of *Slp1*

To check transcription of *Slp1* in immature leaf, mature leaf and immature inflorescence, RNA was isolated from these tissues and cDNA was synthesized using 1 μg of total RNA after on column-DNase treatment (Zymoresearch, CA, USA). Equal volume of cDNA was PCR amplified using the primers specific for *Slp1* (Slp1-RT-F and Slp1-RT-R) and ubiquitin-conjugating enzyme (UbCE; Sb09g023560) as internal control (Shakoor *et al*., 2014; Markovich *et al*., 2015). The primer pair used to amplify *UbCE* was UbCE-RT-F and UbCE-RT-R. Nucleotide sequences of primers are given in Supporting Information Table S1.

To measure relative transcript abundance of *Slp1* in ASZ-2, ASZ-1, ASZ, ASZ+1 and the youngest mature leaf, three independent sorghum seedlings with immature leaf length between 13-16 cm were used. The youngest mature leaves were used as a measure of background transcription level of *Slp1* as silica cells in these regions are dead and silicification is no longer taking place in them (Kumar & Elbaum, 2018). RNA isolation and cDNA synthesis were carried out as written above. Reactions were performed using SYBR mix (Invitrogen, MA, USA) in 7300 Real Time PCR machine (Applied Biosystems, MA, USA) as explained before (Markovich *et al*., 2015) except that the primer concentrations were kept at 150 nM each. A melt curve analysis at the end of the PCR cycle was carried out to ensure that only one PCR product was formed. *Slp1* and *UbCE* transcripts were amplified using primers as mentioned above. In addition, we also amplified transcripts of an RNA recognition motif-containing protein (*RRM;* Sb07g027950) as another internal control (Shakoor *et al*., 2014) using the primers RRM-RT-F and RRM-RT-R (Table S1).

Using *UbCE* as internal control, the relative transcript abundance of *Slp1* was calculated according to the formula of (Pfaffl, 2001). The relative transcript level of *RRM* is also reported, showing that the transcript level of the two house-keeping genes do not change significantly in the tested tissues (Supporting Information Fig. S1).

### Immunolocalization of Slp1 in sorghum leaves

Leaf tissues from ASZ and mature leaves were cut into about 1 mm X 1 mm pieces and fixed in 4% para-formaldehyde (w/v) containing 0.05% triton X-100 using vacuum infiltration. Blocking was done in PBS buffer containing 0.1% BSA. The tissue sections were incubated with purified polyclonal antibody against Peptide-1 (10 μg ml^−1^) for one hour at room temperature. After washing, the tissue sections were incubated with secondary antibody (goat anti-rabbit IgG) tagged with Alexa flour 488. The tissue sections were further washed and incubated for 10 min in propidium iodide (5 μg ml^−1^) solution to stain the cell wall red. Tissue sections undergone the same treatment (i) with pre-immune serum, or (ii) without the primary or the secondary; or without both the antibodies worked as control. The tissue segments were mounted on microscopic slide and observed under Leica SP8 inverted confocal laser scanning microscope with solid-state laser (Wetzlar, Germany). The excitation wavelength was 488 nm while the emission filter range was 500-550 nm for Alexa flour 488. Propidium iodide was excited at 514 nm and the emission filter range was 598-634 nm.

### Transient overexpression of Slp1 in tobacco (*Nicotiana benthamiana* Domin)

pBIN-19 plasmid was modified by inserting CaMV35S promoter in between HindIII and SalI restriction sites, whereas GFP followed by NOS terminator was inserted in between XbaI and EcoRI restriction sites. The resulting plasmid (pBIN-GFP) was used as control plasmid to drive the expression the GFP under the control of CaMV35S promoter. Primers Slp1-SalI-F and Slp1-XbaI-R (Table S1) were used to amplify the full length of Slp1 without the stop codon and flanked with the restriction sites SalI and XbaI, respectively. The PCR product was digested with SalI and XbaI and ligated to the SalI and XbaI digested pBIN-GFP plasmid. The expression cassette consisted of CaMV35S promoter-Slp1-GFP-NoS terminator. We immobilized the plasmids into *Agrobacterium tumefaciens* (strain EHA105) and infiltrated *Nicotiana benthamiana* source leaves on the abaxial side with slight modifications from the reported protocol (Li, 2011). Our resuspension solution consisted of 50 mM MES buffer, 2 mM NaH_2_PO_4_ supplemented with 200 μM acetosyringone. After 48 hours, leaf pieces were observed under laser scanning confocal microscope (excitation: 488 nm; emission filter range: 500-550 nm).

### Transient overexpression of Slp1 in sorghum

We created a construct exploiting the maize dwarf mosaic virus (MDMV) genome (https://patentscope.wipo.int/search/en/detail.jsf?docId=WO2016125143). We fused the expression cassette containing a CaMV35S promoter driving the expression of Slp1 and *nos*-terminator in between *Age*I and *Apa*I restriction sites to the MDMV genome as follows. We amplified full length of Slp1 except the start and stop codons using the primers Slp1-AgeI-F and Slp1-ApaI-R (Table S1). After digestion of the PCR product with AgeI and ApaI restriction enzymes, we ligated the digested PCR product with AgeI and ApaI digested MDMV-GUS plasmid. The resulting plasmid was named MDMV-Slp1. The plasmids were coated on gold microparticles (1 μm diameter, Bio-Rad, California, USA) and bombarded on about one week old sorghum seedlings according to the protocol of (Jose-Estanyol, 2013) with the differences explained below. Sorghum seeds were surface sterilized, germinated and then grown with their roots immersed in tap water for up to seven days before they were bombarded. Just before bombardment, the seedling roots were quickly taken out of water, and a bunch of five seedlings were arranged and loosely stuck in the center of a petri dish and bombarded with the plasmid coated gold particles using 1100 psi rupture discs. After bombardment, the seedlings roots were wrapped in moist tissue paper and put in a beaker with tap water for 24 hours, after which the bombarded seedlings were transferred to soil. The seedlings were grown for about three weeks and the phenotypes in the leaves showing viral symptoms were observed in the SEM. RNA was isolated from the young leaves of the infected plants and cDNA was synthesized. Primers derived from the MDMV-Slp1 plasmid (MDMV-Slp1-F and MDMV-Slp1-R, Table S1) were used to amplify Slp1 with viral flanking regions, or primers from viral protein coat (MDMV-Coat-F and MDMV-Coat-R, Table S1) were used to ascertain the viral infection in the control plants (infected by MDMV-GUS plasmid). We conducted this experiment twice, each time starting with 15 independent biological replicates and analysing altogether 10 plants showing strong viral symptoms.

## Results

### Screening for putative silicification protein(s)

In order to find candidate silicification regulating factors, we analysed a publicly available data set which reports the influence of silicon treatments on gene expression in wheat (GEO NCBI dataset GSE12936) (Chain *et al*., 2009), as both sorghum and wheat belong to Poaceae (grass) family. We searched for genes that co-express with wheat silicon transporters (*TaLsi1, TaLsi2, TaLsi6*) irrespective of the plant pathogenic infection status, using Comparative Co-Expression Network Construction and Visualization (CoExpNetViz) online tool (Tzfadia *et al*., 2016). Once co-expressed genes were identified, their probe sequences were extracted and used to BLAST against the sorghum genome. The BLAST yielded a list of 18 sorghum orthologues genes (e-value > 100 was used as orthology cutoff) (Table S2). The primary sequence of all the initially screened proteins were analysed for the theoretical isoelectric point (pI) using ProtParam tool in ExPASy server (https://web.expasy.org/protparam/); and the presence of predicted signal peptide (SignalP server). Since positively charged amino acids have been shown involved in biological silicification, we discarded all the proteins with predicted pI value less than 7. We also rejected the proteins that lacked signal peptide as a silicifying protein must be secreted outside the silica cell membrane in order for silicification to take place in the paramural space. Proteins were further screened for the presence of internal repeat units as found in several silica precipitating proteins (Kröger *et al*., 1999; Kauss *et al*., 2003; Shimizu *et al*., 2015), abundance of histidine, glutamic acid (Shimizu *et al*., 2015), lysine (Kröger *et al*., 1999; Kauss *et al*., 2003) or serine, glycine-rich motifs, and frequency of proline-lysine and proline-glutamic acid residues (Harrison, 1996). Based on our selection criteria, we identified a protein in sorghum (Sb01g025970) that had seven repeat units rich in lysine, similar to silaffins from *Cylindrotheca fusiformis*, a diatom species (Kröger *et al*., 1999); histidine-aspartic acid rich regions similar to glassin from a marine sponge, *Euplectella* (Shimizu *et al*., 2015), and several proline residues as in a proline-rich protein (PRP1) from cucumber (Kauss *et al*., 2003). We functionally characterized this protein and named it Siliplant1 (Slp1).

### Molecular architecture of Sorghum Slp1

Sorghum Slp1 comprises of 524 amino acids with a predicted N-terminal signal sequence (Fig. 1). The signal sequence is predictably cleaved between amino acid positions 24 and 25. Further sequence characterization using TargetP predicted Slp1 to be a secretory protein with high probability. BLAST-based homology search revealed that Slp1 is a single copy gene in sorghum and belongs to a family of proline-rich proteins with unknown function. The transcript of this protein is highly abundant in young leaves and inflorescences (MOROKOSHI, © 2018 RIKEN, query Sobic.001G265900). Slp1 is rich in proline, lysine, and histidine, respectively comprising 20%, 13%, and 11% of total amino acids. The theoretical pI value of the protein is 9.28. Structure predictions using GOR4 (Combet *et al*., 2000) suggest that about 80% of the protein is random coil and 14% is alpha helix. IUPred suggested that Slp1 has intrinsically disordered tertiary structure (Dosztányi *et al*., 2005a,b). Slp1 has repeat units (R1 to R7) rich in proline, lysine, and glutamic acid (P, K, E-rich domain) with the consensus sequence KKPXPXKPKPXPKPXPXPX. Near a wide range of physiological pH, close to half of this domain is positively charged (lysine-rich) and the rest of the domain is negatively charged (aspartic acid-rich). Five of the repeat units (except R1 and R3) have a histidine and aspartic acid (H, D) rich domain with the consensus sequence DXFHKXHDYHXXXXHFH immediately preceding the P, K, E-rich domain. This aspartic acid rich domain is negatively charged under physiological pH. In five repeats (except R2 and R4) there is a third domain following the P, K, E-rich domain, rich in proline, threonine and tyrosine (P,T,Y-rich) with the consensus sequence YHXPXPTYXSPTPIYHPPX. Before the start of the repeat units (amino acids 1-148), there is a region rich in alanine, serine, glycine and valine (comprising together 45% of the region). At the end of R1, R3, and R5, we found an RXL domain. This domain acts as recognition site for unknown proteases that cleave at the C-terminus of the leucine residue of the RXL domain in many bio-silica associated proteins in diatoms (Kröger *et al*., 1999; Wenzl *et al*., 2008; Scheffel *et al*., 2011; Kotzsch *et al*., 2017). Despite these common features, Slp1 does not show sequence similarity with any protein known to be involved in bio-mineralization.

**Fig. 1.**
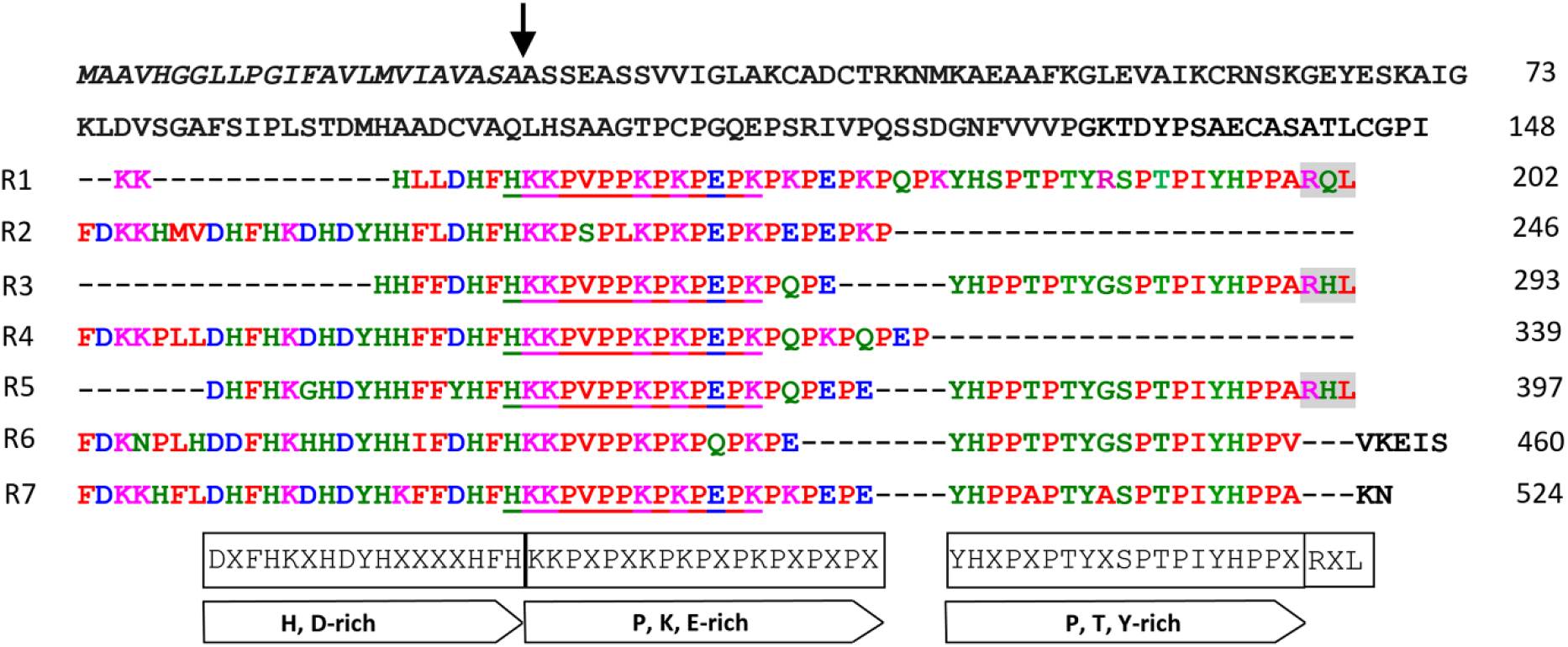
Primary sequence of *Sorghum bicolor* Slp1, showing the N-terminal signal sequence and the seven repeats (R1-R7). The predicted signal sequence is shown in italics. The arrow denotes the predicted signal sequence cleavage site. All the repeats (R1-R7) have a proline, lysine and glutamic acid rich (P, K, E-rich) domain forming a zwitterionic center. In addition, a histidine-aspartic acid rich (H, D-rich) domain, negatively charged near physiological pH precedes the P, K, E-rich domain in repeats 2, 4, 5, 6 and 7. Whereas, a proline, threonine and tyrosine rich (P, T, Y-rich) domain follows the P, K, E-rich domain in repeats 1, 3, 5, 6 and 7. At the end of repeats 1, 3 and 5, there is an RXL domain (shaded) which serves as cleavage site for unknown proteases in many bio-silica associated proteins in diatoms. The underlined sequence is Peptide-1 which appears five times in the primary sequence and was used for raising antibodies and for silica precipitation assays. The consensus sequence for the domains is given in boxes below each domain, where X denotes any amino acid.

### Silica precipitated *in vitro* by Peptide-1 derived from Slp1 sequence

In order to test the possible activity of Slp1 as a silica mineralizer we synthesized the peptide sequence HKKPVPPKPKPEPK (Peptide-1) which appears 5 times in the P, K, E-rich domain of Slp1 primary sequence. When metastable (90.9 mM) silicic acid solution was added to Peptide-1 solution (final peptide concentration 1.82 or 1.36 mg ml^−1^) at neutral pH, the reaction mixture became cloudy within seconds, and very fine, white sand-like powder precipitated. In contrast, a mutant peptide, where the lysine residues in Peptide-1 were replaced by alanine (HAAPVPPAPAPEPA; Peptide-3), did not precipitate powder-like silica at 90.9 mM silicic acid. Gel-like material formed when the reaction mixture was centrifuged (Fig 2a). Scanning electron micrographs of the powder sediment, forming within 5 minutes revealed spheres of about 0.5 microns in diameter (Fig. 2b). Silica was also precipitated by Peptide-1 (1.36 mg ml^−1^) at silicic acid concentration of 5 mM (pH=7.0), which is in the lower concentration range found in grass leaves (Casey *et al*., 2003). The precipitation was invisible to the naked eyes even after 30 minutes of incubation. However, SEM examination of the sediment showed that there indeed was silica precipitation and the diameter of an individual silica nanosphere was about 250 nm (Fig. 2c). In control solutions (without peptide) at 90.9 and 5 mM silicic acid, precipitation was not observed by SEM. High resolution transmission electron micrographs (HRTEM) of particles of 0.5 microns revealed molecular scale dark and bright regions, possibly resulting from the contrasting electron density of the peptide and poly-siloxane chains. The dark and bright patterns create a short range periodic order a few nanometers long and 1 nm thick.

**Fig. 2.**
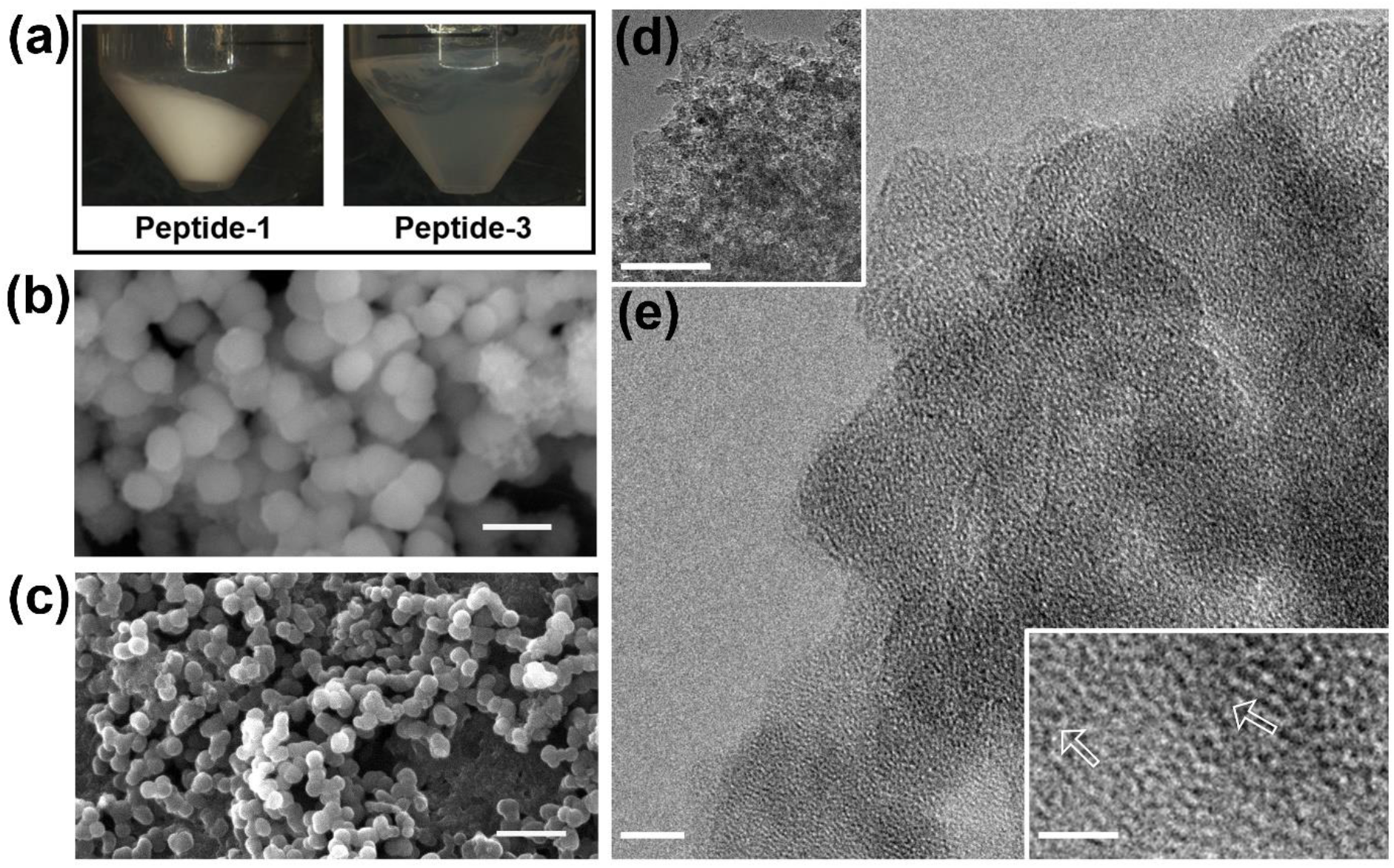
Imaging of silica precipitation by peptides derived from *Sorghum bicolor* Slp1 sequence. (a) Sand-like powder sediment produced from metastable (90.9 mM) silicic acid solution with Peptide-1, while a gel-like material formed after several rounds of centrifugation of silicic acid solution with Peptide-3. (b) Scanning electron micrograph of the powder sediment formed by Peptide-1 at 90.9 mM silicic acid. (c) Scanning electron micrograph of the silica sediment formed by Peptide-1 at 5 mM silicic acid. (d) High-resolution transmission electron microscopy (HRTEM) of one particle imaged in panel (b) reveals a mesoporous structure on a nanometric scale. Bar, 5 nm. (e) Dark and bright granulation have short-range order, as marked by arrows in the inset panel. Bars of 1 μm in panels (b, c), 60 nm μm in (d), 5 nm in (e), and 2 nm in (e-inset).

### Raman and NMR spectroscopic characterization of Peptide-1 – silica precipitate

The precipitated silica with Peptide-1 at 90.9 mM silicic acid was characterized by Raman and magic angle spinning (MAS) solid-state nuclear magnetic resonance (ss-NMR) spectroscopies (Fig. 3). Raman spectrum of the sediment showed that the mineral is silica, recognized by the peaks at 489 cm^−1^ and 997 cm^−1^ (Aksan *et al*., 2016). These peaks are missing in the Raman spectrum of Peptide-1 (Fig. 3a). Further analysis indicated chemical interactions between the silica surface and the amino group of the lysine side chain (Supporting Information Notes S1).

**Fig. 3.**
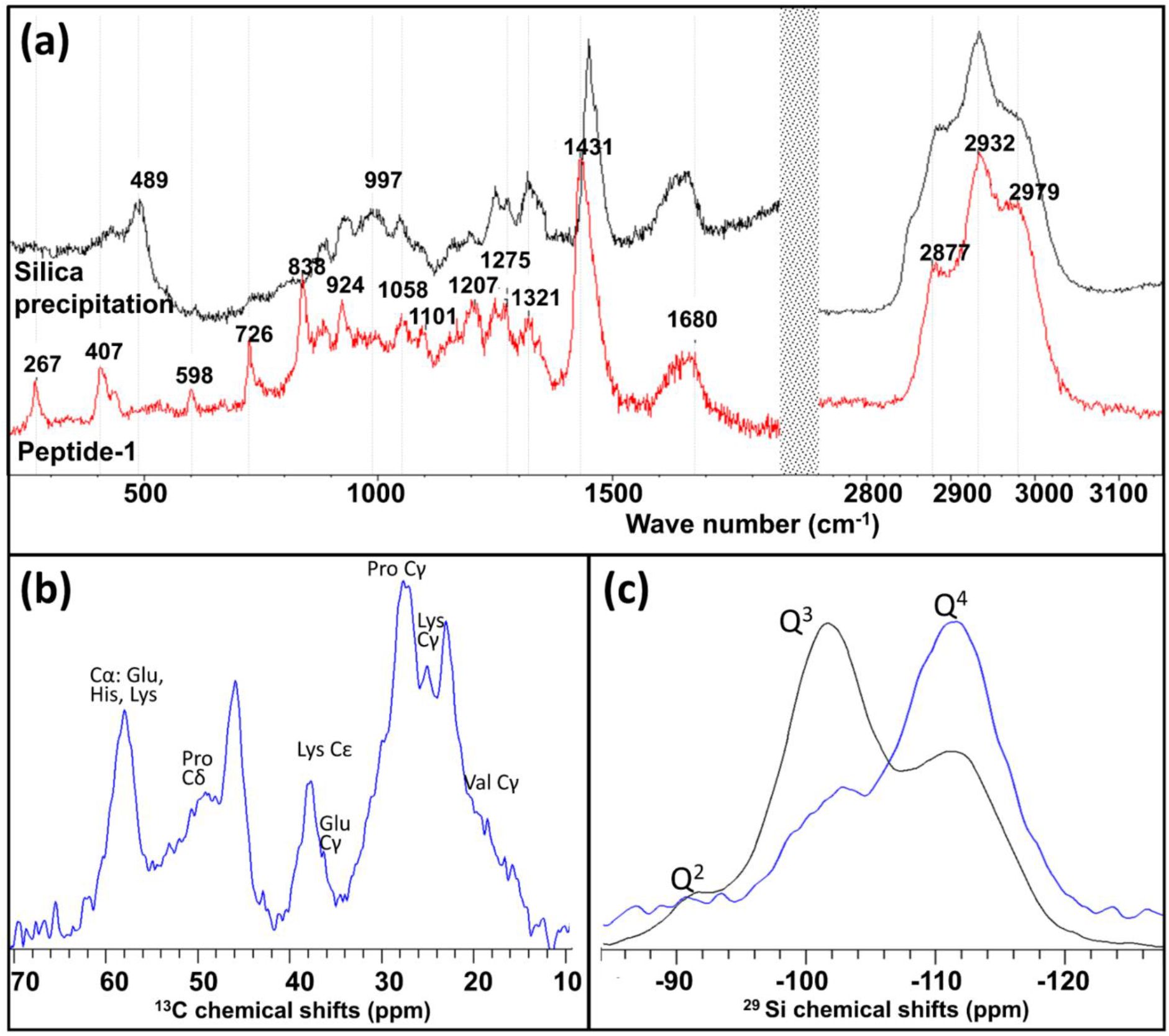
Characterization of the peptide-silica sediment using Raman and NMR spectroscopy. (a) Raman spectra of Peptide-1 (red line) and the sediment formed with Peptide-1 (black line). The spectra show that the mineral is silica, with altered surface, and that the peptide is bound to the mineral through the lysine aliphatic amine and proline ring residues and the -COO^-^ C-terminal group. See Note S1 for peak assignment. (b) Spectrum of the peptide-silica sediment, measured by magic angle spinning (MAS) solid-state nuclear magnetic resonance (ss-NMR) ^1^H-^13^C cross polarization. The NMR signals, typical to aliphatic bonds in the amino acid sidechain, are shifted to a high field by about 2 ppm. These shifts reflect a shielding effect of the silica, suggesting that the sidechains are bound to the mineral. (c) Spectra of ^29^Si showing peaks of Si-(OSi)_2_(OH)_2_ (Q2); Si-(OSi)_3_(OH) (Q3); and Si-(OSi)_4_ (Q4). Direct polarization (black line) samples all the Si atoms in the sample, while ^1^H-cross polarization (blue line) samples Si atoms in proximity to protons.

By conducting large scale silica precipitation assay with Peptide-1 at 90.9 mM silicic acid, we could collect 113 mg of dried silica, sufficient for NMR analyses. Confirmation of the HRTEM and Raman findings that the peptide was complexed with the silica was given through NMR ^1^H-^13^C cross polarization measurements (Fig. 3b). The spectrum shows narrow peaks (full width at half maximum (FWHM) was 282 Hz), as compared with spectra of a diatom peptide complexed with silica (FWHM of 489 Hz) (Geiger *et al*., 2016). The narrow peaks allowed us to identify many of the amino acid sidechain carbons, with peaks shifted to a higher field by about 2 ppm. Such shifts, caused by the silica shielding the magnetic field that is felt by the peptide, indicate of close proximity of the mineral to the peptide.

In order to study the structure of the mineral we collected NMR direct polarization signal from Si atoms (Fig. 3c). ^29^Si NMR can detect the number of groups of -OSi and -OH bound to a central Si atom. A Qn band is defined as Si-(OSi)_n_OH_4-n_ (Engelhardt & Michel, 1987). The relative intensities of the Q4:Q3:Q2 peaks in the spectrum were 62:33:5, indicating that there are about 5 bulk (Q4) Si atoms for every 3 surface (Q3+Q2) atoms. To examine further the silicon atoms at the surface we excited them through near-by hydrogen atoms, by cross polarization measurement (Fig. 3c). This measurement revealed that in addition to surface Q3 and Q2 we detected some Q4 siloxane species bound to surface oxygen. These Si atoms were excited by protons from nearby silanols and water, indicating their closeness to the surface of the silica particle.

### Expression pattern and subcellular localization of Slp1

Our sequence-based predictions and *in vitro* results suggested a role for Slp1 in silica deposition. To investigate whether the expression profiles of Slp1 correlate with silica deposition times and locations, we raised an antibody against Peptide-1. Using Western hybridization, we detected several bands in immature leaves and inflorescence. Slp1 expression was not detected in roots and mature leaves (Fig. 4a). In correlation with the protein expression profile, we detected RNA transcription of *Slp1* in immature leaves and inflorescences, but not in mature leaves (Fig. 4b). This profile matches regions of active silica deposition, which occurs only in young leaves but not in mature leaves. RNA transcript profile of *Slp1* along immature, silicifying leaf tissues showed that *Slp1* is strongly transcribed in the leaf base, reaching the highest levels just before the active silicification zone (ASZ-1). The transcription levels dropped by a factor of about 15 in the ASZ, and further reduced in older tissues of immature leaves (Fig. 4c). In the youngest mature leaf tissue, we found that *Slp1* is transcribed at a basic, low level constituting the background transcription level in these leaves. Background transcription level was about 1/19,000 that of the maximal transcript in ASZ-1.

**Fig. 4.**
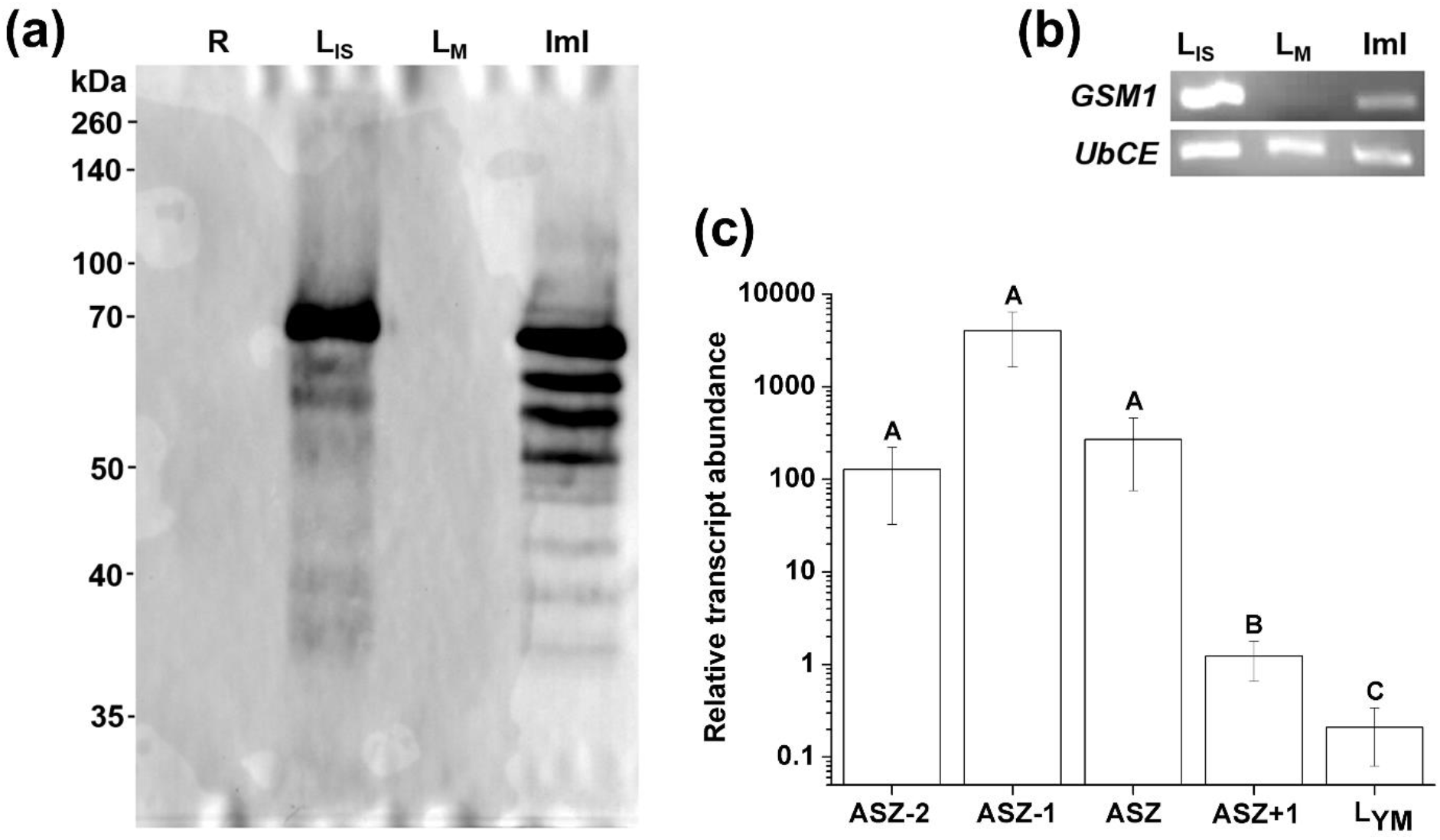
Expression profile of *Sorghum bicolor* Slp1. (a) Western blot of crude protein extract from roots (R), immature silicifying leaves (L_IS_), mature leaves (L_M_), and immature inflorescences (ImI), detected by an antibody against Peptide-1. Slp1 was expressed in developing leaves and inflorescence, but not in roots and mature leaves. Multiple sized bands in the inflorescence may indicate that Slp1 is processed differently in immature leaf and inflorescence. (b) RNA transcription of *Slp1* in immature silicifying leaves (L_IS_), mature leaves (L_M_), and immature inflorescences (ImI). In accordance with protein translation, transcription was detected only in the immature tissues. *UbCE* (ubiquitin-conjugating enzyme) was used as internal control gene. (c) RNA transcript profile of *Slp1* along immature silicifying leaf tissues. Maximal transcript was found in ASZ-1, which lies just below the Active Silicification Zone (ASZ). Error bars indicate standard deviation (n=3).

To further correlate Slp1 with silica deposition, we tested its distribution in leaf epidermal cells using antibody against Peptide-1. In the ASZ, the antibody bound to silica cells (Fig. 5a,b). We noticed the antibody signal in either the cytoplasmic volume or near the cell periphery of silica cells (Video S1). Further image analysis suggested that the cytoplasmic immunofluorescence signal originated from vesicles (Fig. 5c). In mature leaves, where silica cells are silicified and dead, the antibody hybridized only to the boundary of silica cells (Fig. 5d,e). No vesicles were identified by image analysis (Fig. 5f). Immunolocalization with the pre-immune serum, or the reactions lacking any one or both the antibodies as control did not fluoresce (Fig. S2).

**Fig. 5.**
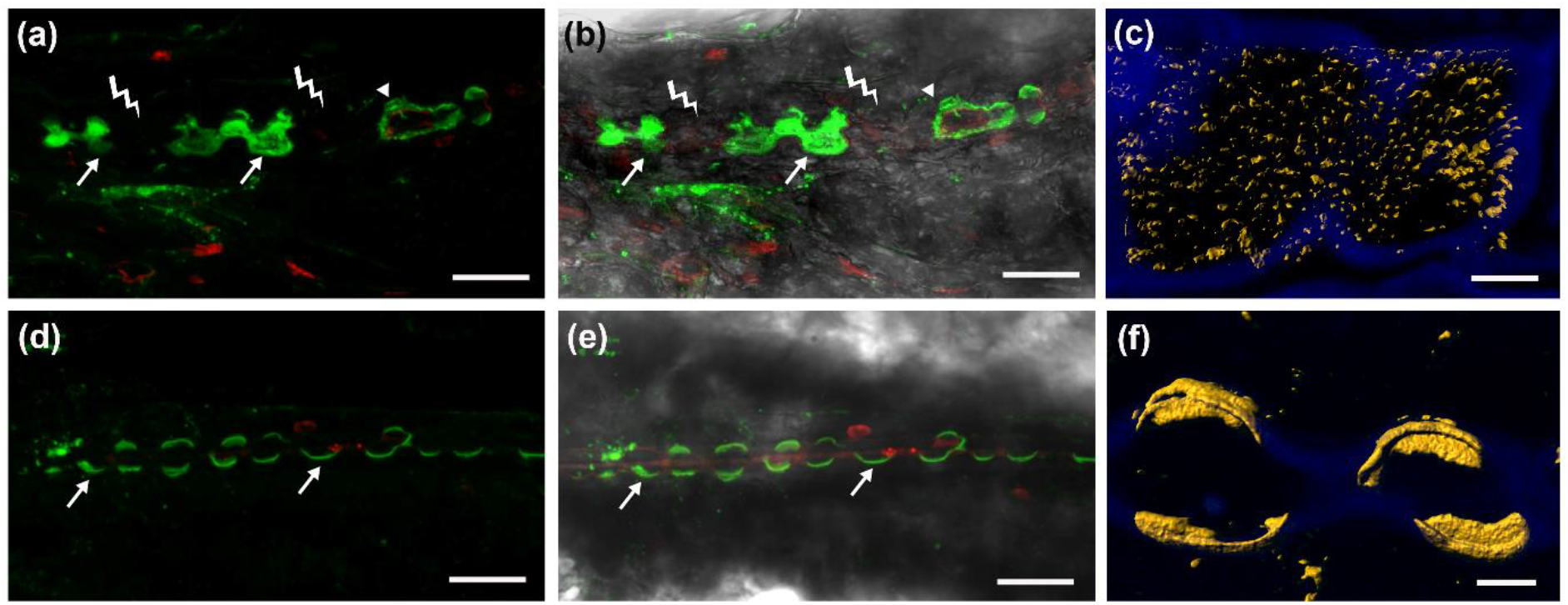
Immunolocalization of *Sorghum bicolor* Slp1 in the active silicification zone (ASZ) of young leaves (a-c), and near the tips of fully silicified mature leaves (d-f). In sorghum leaves, silica cells usually pair with cork cells (indicated by thunder sign) and exist as long chains of alternating silica and cork cells; but near the leaf tip, cork cells are absent and silica cells fuse to form long polylobate bodies. (a) Immunolocalization (green fluorescence) shows that Slp1 is localized to silica cells of sorghum leaf active silicification zone (ASZ), inside the cytoplasm (arrows) or near the cell periphery (arrowhead). (b) Fluorescence image in (a) merged with the corresponding brightfield image. (c) The anti-peptide-1 antibody fluorescence was processed to select for punctuated regions (pseudocoloured yellow) in the ASZ. The Slp1 appears in packets in the cytoplasm and cell boundary. (d) The anti-peptide-1 antibody (green fluorescence) binds to the edges of silicified silica cells (arrows) in mature leaves. (e) Fluorescence image in (d) merged with the corresponding brightfield image. (f) Processed fluorescence of mature leaf image demonstrates that peptide-1 (pseudocoloured yellow) is embedded inside the polymerized silica. Red fluorescence is from propidium iodide, blue is background autofluorescence of the cell walls. Bar 25 μm in panels (a, b, d, e) and 4 μm in panels (c) and (f).

The presence of N-terminal signal peptide in Slp1 predicted it to be secretory in nature. Hence, to verify this prediction, we made translational fusion of green fluorescent protein (GFP) to the C-terminal of full length Slp1 and transiently overexpressed it in tobacco. Similar to our observations in the silica cells of the ASZ, the cells expressing the fusion protein fluoresced in discrete packets distributed throughout the cytoplasm (Fig. 6a,b). In addition, we could identify packets fusing to the cell membrane as well as diffused fluorescence at cell boundaries (Fig. 6c,d, Video S2). In control tobacco plants that overexpressed GFP without Slp1, the fluorescence was uniform in the cytoplasm and nucleus, and lower between cells (Fig. 6e,f). Mock infiltration of tobacco leaves using MES buffer did not fluoresce (Fig. 6g,h). Thus, our results show that native Slp1 is packed in vesicles inside silica cells and suggest that silica cells release Slp1 packages to the paramural space. Since the apoplasm of grass leaves is super-saturated with silicic acid, secretion of the Slp1 to the apoplast may lead to immediate silica precipitation.

**Fig. 6.**
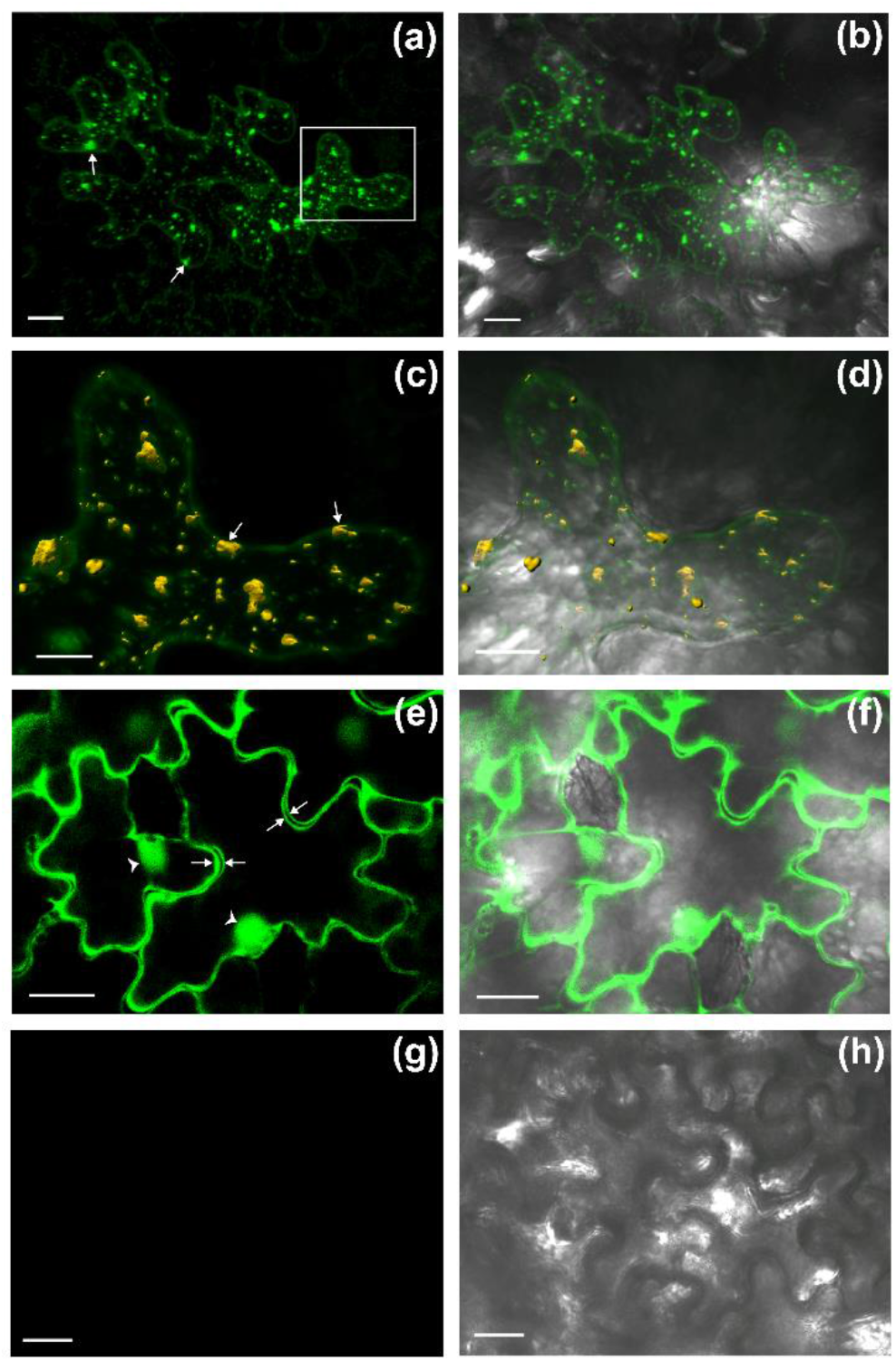
Fluorescence confocal microscopy showing the secretory nature of *Sorghum bicolor* Slp1. (a) Slp1 fused with GFP was transiently overexpressed in tobacco leaf using *Agrobacterium tumefaciens*. The green fluorescence, marking the location of Sil, was found in packets inside the cytoplasm (arrows), while diffused green fluorescence can also be seen along the margin of the cell. White rectangle marks a region enlarged in panel (c). (b) Fluorescent image in (a) merged with the corresponding brightfield image. (c) Segmenting the green fluorescence to diffuse (green) and punctate (pseudocoloured yellow) regions shows packets fusing to the cell membrane (arrows) as well as diffused fluorescence at cell boundaries. (d) Fluorescent image in (c) merged with the corresponding brightfield image (e) Control plants expressing GFP without Slp1 showing green uniform fluorescence of the cytoplasm and nucleus (arrowheads). Arrows indicate the cytoplasm of adjacent cells, demonstrating the low fluorescence between cells. (f) Fluorescent image in (e) merged with the corresponding brightfield image. (g) Mock infiltration of tobacco leaves using MES buffer did not fluoresce. (h) Fluorescent image in (h) merged with corresponding brightfield image. Bars in panels (a, b, e, f, g, h) are 20 μm, and in (c, d) are 10 μm.

### Functional characterization of Slp1 *in planta*

To elucidate the role of Slp1 *in planta*, we transiently overexpressed it in sorghum, using a construct derived from maize dwarf mosaic virus. Compared with the wild type plants, control sorghum plants infected with virus lacking the Slp1 sequence showed infection lesions and viral symptoms (Fig. S3) but no unusual silica deposition (Fig. 7a-f). In contrast, the Slp1 overexpressing plants had ectopic silica deposition in patches, especially close to the veins and where viral symptoms were observed (Fig. 7g-i). Silica was also deposited in cells which are not usually silicified in sorghum leaves, like guard cells, cork cells and long cells (Fig. 7j-l). SEM in the back-scattered electron mode suggested the silicification intensity to be of a level similar to that in silica cells. Similar observations were recorded in ten independent biological replicates. Energy-dispersive X-Ray spectroscopic signal for silicon from the heavily silicified patches in the Slp1 overexpressing leaves was much higher than in the wild type and control leaves. In correlation, the locations of high Si contained also high concentrations of oxygen, low concentrations of carbon, and similar background levels for other elements.

**Fig. 7.**
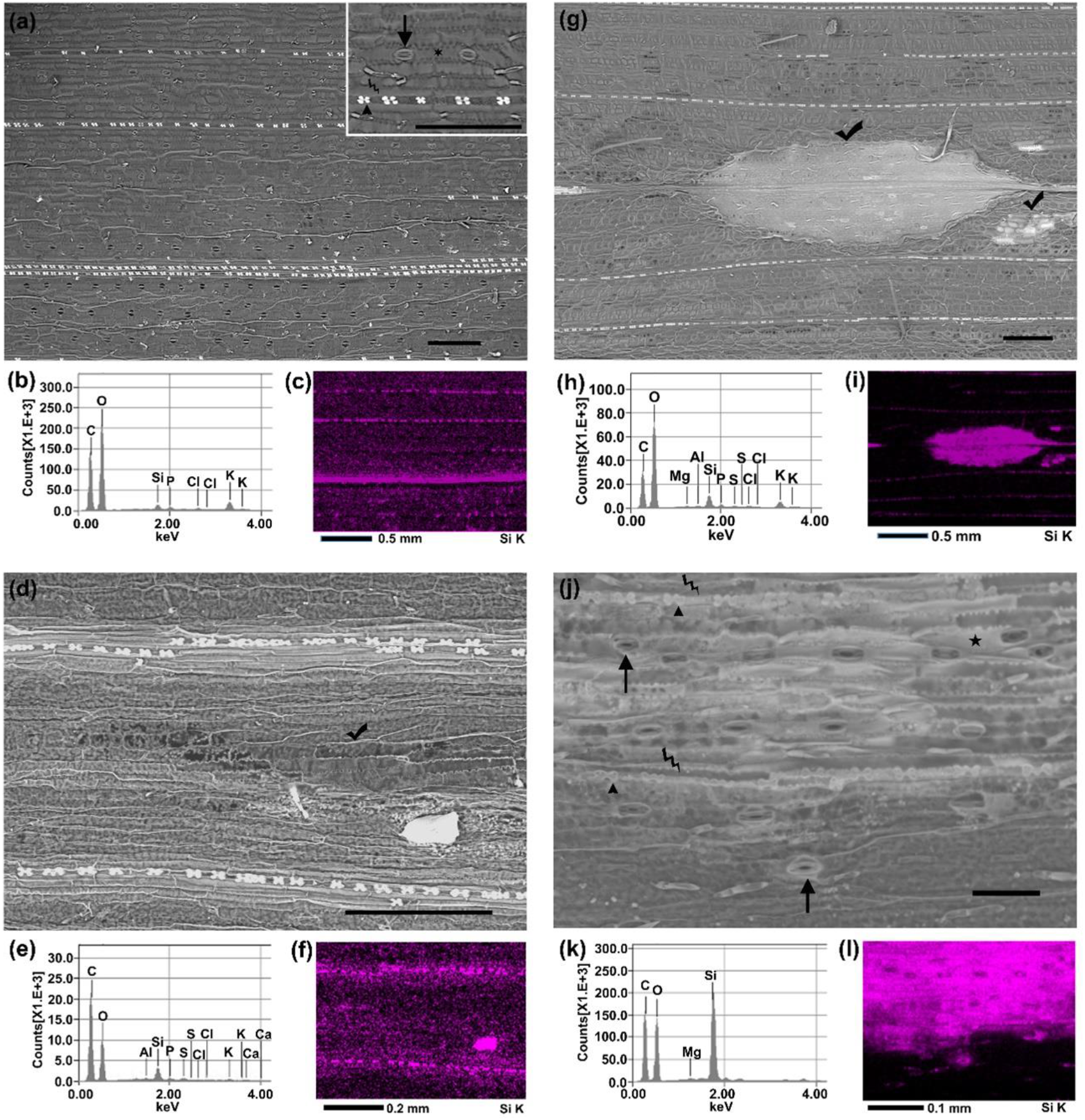
Overexpression of *Sorghum bicolor* Slp1 in sorghum. Arrowheads indicate silica cells, arrows- stomata, thunder signs- cork cells, stars- long cellsand tick mark indicate viral lesions. (a,d,g,j) Scanning electron microscope (SEM) image in back-scattered electron (BSE) mode. Silicified cells appear brighter compared with the background. (b,e,h,k) Energy-dispersive X-ray spectra (EDS) of the corresponding SEM image showing the comparative elemental composition of the scanned area. (c,f,i,l) EDS map for Si-signal form the corresponding SEM image. Maps are pseudocoloured. Black colour means no signal and increasing intensity of the colour denotes higher density of Si atoms. (a) Wild type sorghum mature leaf scanned by SEM under low magnification showing the general silicification pattern in a mature leaf. Inset: epidermal cell types are indicated in a higher magnification image. (b) EDS of the image in (a). (c) EDS map for Si-signal of the image in (a). (d) SEM image of control plants showing silicification only at usual locations. Dust particle with high Si content is seen on the bottom right. (f) EDS map for Si of the image in (d). (g) SEM image of ectopic silica deposition in a viral lesion in Slp1 overexpressing plant. (h) EDS of the image in (g). (i) EDS map for Si of the image in (g). (j) SEM image of Slp1 overexpressing plants showing high intensity silica deposition in cells that do not usually silicify in wild type plants. All epidermal cell types can be seen silicified. (k) EDS of the image in (j). (l) EDS map for Si of the image in (j). Bars represent 200 μm in panels (a; d and g); and 50 μm in panel (j).

## Discussion

Silicification in plants may be passive or biologically controlled depending upon the site of deposition (Kumar *et al*., 2017b). Out of several cell types that deposit silica in grasses, silica cells are unique as more than 95% of cells silicify in young, elongating leaves. Their mineralization requires metabolic activity and is independent of transpiration (Sangster and Parry 1971). The fast silica precipitation over a few hours (Lawton, 1980) in addition to the formation of the mineral at the cell wall, as opposed to the cytoplasm (Kumar *et al*., 2017a; Kumar & Elbaum, 2018), point to a factor exported from the cells that induces biogenic silica formation. Our work shows that silica cells express and export the protein Slp1 to the apoplast, timed with silicification in the paramural space (Kumar *et al*., 2017a; Kumar & Elbaum, 2018). It is obvious that silica deposition depends on the presence of silicic acid which is absorbed by roots from the soil. Typical silicic acid concentrations measured in the xylem sap are 5-7.5 mM for wheat (Casey *et al*., 2003), 5-25 mM for rice (Ma *et al*., 2002; Mitani & Ma, 2005), and 4, 7-12 mM in sorghum seedlings and mature plants, respectively (our unpublished data). Since Slp1 overexpression caused deposition in all epidermis cell types, we can conclude that it actively precipitates silica at *in planta* silicic acid concentrations. Furthermore, the fact the silica formed ectopically in stomata and long cells indicates that Slp1 is sufficient to cause precipitation in those cells. Based on our results we conclude that Slp1 in its native form actively deposits silica *in planta*.

### Expression patterns are in correlation with silica deposition in silica cells

Slp1 expression was not detected in roots where silica cells do not form. Furthermore, we could not detect Slp1 expression in mature leaf tissues, in correlation with the absence of viable, active silica cells (Kumar & Elbaum, 2018). This suggests that silica deposition in live silica cells is dependent on Slp1, while silica deposition in cell walls and possibly other locations is governed by other means such as specific cell wall polymers (Fry *et al*., 2008; Law & Exley, 2011; Kido *et al*., 2015; Brugiére & Exley, 2017; Kulich *et al*., 2018). Our report on Slp1 expression in immature inflorescence is consistent with published transcriptomic data, showing *Slp1* transcript both before and after inflorescence emergence (Table S5 of Davidson et al. 2012). We propose that some cell types in the inflorescence employ Slp1 to produce silica without transpiration. These may be glume abaxial epidermal cells that silicify before inflorescence emergence and macro-hairs and the long cells that are located on the abaxial epidermis of lemma, which lacks stomata and thus the transpiration stream (Hodson *et al*., 1984; Hodson, 2016; Kumar *et al*., 2017b).

### Possible post-translational modifications

Some indication for Slp1 post-translational modifications may be the time gap between its highest transcription level in the ASZ-1 tissue, and the highest silicification activity in the ASZ (Kumar *et al*., 2017a; Kumar & Elbaum, 2018). One such possible modification may be glycosylation (Elbaum *et al*., 2009). Processed Slp1 molecules are then packed inside vesicles (Lawton, 1980) and stored in the cytoplasm for later release (Alberts *et al*., 2002) at the time of silicification. Modifications may be tissue and species specific. The difference in the size of the Western hybridization signal between immature leaf and inflorescence that we observed may be attributed to differential processing of Slp1 in these two tissues. Slp1 has three RXL domains in the primary structure. RXL domains are proteolytically cleaved in many diatom biosilica associated proteins (Kröger *et al*., 1999; Wenzl *et al*., 2008; Scheffel *et al*., 2011; Kotzsch *et al*., 2016, 2017). It would be interesting to study if Slp1 undergoes alternative processing, similar to *Thalassiosira pseudonana* (a diatom species) silaffin (Poulsen & Kröger, 2004).

### Slp1 precipitates together with silicic acid creating highly porous biosilica

Binding of anti-Peptide-1 antibody in the boundary of dead silica cells of mature leaves suggests that as Slp1 templates the silica precipitation, it is caught inside the silica structure. Silicanin-1, a biosilica associated protein from *T. pseudonana* gets embedded inside biosilica structure and is accessible to anti-silicanin-1 antibody (Kotzsch *et al*., 2017). Similarly, anti-Peptide-1 antibody may also have access to the epitope remains on the surface of the deposited silica in silica cells. Our *In-vitro* experiments support such entrapment, with significant shifts in the NMR and Raman peaks, indicating close proximity between the peptide and the mineral. The periodic short-range order, as seen in HRTEM, suggests a very intimate interaction of the forming mineral and the peptide, similarly to a peptide derived from Silaffin3, a silica precipitating protein from diatom (Iline-Vul *et al*., 2018). The mineral that we produced *in vitro* contains many surface Si atoms, forming an open mesoporous structure. Similarly, native plant silica may form a permeable matrix that will allow movement of solutes. Diffusion of silicic acid through the mineral is required in order for silica cells to fully silicify, because the mineralization front is at the paramural space (Kumar & Elbaum, 2017, 2018; Kumar *et al*., 2017a).

## Conclusions

Over-expression and localization studies of Slp1 show that this protein is involved in silicification in sorghum leaf silica cells. Slp1 has unique amino acid composition, charge distribution and probably post-translational modifications necessary for its activity. Soon after cell division, silica cells start their preparation for silicification. Slp1 is transcribed, translated, and post-translationally modified, packed inside vesicles and stored in the cytoplasm until the cell is ready to silicify. The vesicles fuse to the membrane and release their content in the paramural space to come in contact with silicic acid. This immediately precipitates open-structured silica that allows the diffusion of more silicic acid from the apoplastic space. The rapid formation of the mineral explains the difficulty associated with finding silicification in an intermediate state. The expression pattern, localization and modifications of Slp1 in the inflorescence bracts need to be studied in detail.

## Accession number

The nucleotide sequence of *Sorghum bicolor* Slp1 has been submitted to NCBI with the GenBank accession number-MH558953. GenBank accession number of the pBIN19 plasmid is U09365.1. The MDMV-GUS plasmid that we used in the current study is covered by a published patent number WO2016125143 (https://patentscope.wipo.int/search/en/detail.jsf?docId=WO2016125143).

## Acknowledgements

S.K. was a recipient of post-doctoral fellowship from Planning and Budgeting Committee (PBC), Council of Higher Education, Israel. This research was funded by the Israel Science Foundation grant 534/14. Partial support to the Centre for Scientific Imaging, Hebrew University (Rehovot campus) by a grant from USAID (AID-ASHA-G 1400005) to the American Friends of The Hebrew University and The Robert H Smith Faculty of Agriculture, Food and Environment is gratefully acknowledged. The authors thank. Shmuel Wolf for the pBIN-GFP plasmid, and Nerya Zexer and Victor M. Rodriguez Zancajo for measuring silicic acid concentration in sorghum sap.

## Author Contributions

S.K. and R.E. planned and designed the research; S.K. identified Slp1 from the list of putative silicification related proteins and performed most of the experiments; N.A.-F. did the NMR spectroscopy and analysed the data with G.G.; S.B. performed parts of the experiments; S.K. and J.A.S.-L. performed immunolocalization experiment under the supervision of Y.H.; O.T. analysed the microarray data and prepared the list of Si responsive genes in sorghum; A.O. prepared the MDMV construct under the supervision of H.V.; S.K. and R.E. analyzed the results and wrote the paper. All the authors commented and approved the final version of the manuscript for publication.

## Supplementary Information

**Fig. S1** Transcript level of *RRM* in relation to *UbCE* as internal control gene, showing that the transcript level of the two house-keeping genes do not change significantly in the tested tissues.

**Fig. S2** Immunolocalization control reactions using pre-immune serum; or lacking either one or both the antibodies.

**Fig. S3** Maize dwarf mosaic virus (MDMV) infected sorghum plants showing symptoms.

**Table S1** List of primers used in the present study.

**Table S2** List of the wheat (genes identified that show significantly differential transcription upon silicon treatment verses the non-treated plants, and their *Sorghum bicolor* homologues.

**Video S1** Confocal microscopy video clip showing Slp1 localization in the cytoplasmic space and near the cell boundary of silica cells.

**Video S2** Confocal microscopy video clip showing the vesicles packed Slp1 fusing to the cell membrane.

**Notes S1** Raman analysis

