## Supplementary Text for "Siliplant1 (Slp1) protein precipitates silica in sorghum silica cells"

**Article acceptance date:** Click here to enter a date.

The following Supporting Information is available for this article:

**Note S1** Raman analysis

**Fig. S1** Transcript level of *RRM* in relation to *UbCE* as internal control gene, showing that the transcript level of the two house-keeping genes do not change significantly in the tested tissues.

**Fig. S2** Immunolocalization control reactions using pre-immune serum; or lacking either one or both the antibodies.

**Fig. S3** Maize dwarf mosaic virus (MDMV) infected sorghum plants showing symptoms.

**Table S1** List of primers used in the present study.

**Table S2** List of the sorghum genes identified by BLAST to homologous genes in wheat that show significantly differential transcription upon silicon treatment verses the non-treated plants.

**Video/Movie S1** Confocal microscopy video clip showing Slp1 localization in the cytoplasmic space and near the cell boundary of silica cells.

**Video/Movie S2** Confocal microscopy video clip showing the vesicles packed Slp1 fusing to the cell membrane.

**Note S1: Analysis of the Raman spectra in Figure 2a**

The Raman analysis compared the spectrum of dried powders of Peptide-1 and the precipitant forming when 90.90 mM silicic acid solution was reacted with 1.36 mg $ml^{-1}$ Peptide-1 solution (Figure 3a). Clear bulk Si-O-Si broad peak is observed at 492 $cm^{-1}$ in agreement with literature values (Aksan *et al.*, 2012). Nonetheless, the surface silanol peak at 997 $cm^{-1}$, is shifted from published values (988 $cm^{-1}$, reported by Aksan *et al.* (2012), or 973 $cm^{-1}$, reported by Gierlinger *et al.* (2008)), supporting surface modifications of the mineral.

A shift in the broad peak of Peptide-1 at 1680 $cm^{-1}$ to lower frequencies indicates a change in the peptide backbone. Such shift may indicate that a random coil structure in the pure peptide is more $\alpha$-helix-like in the sediment (Anthony, 1982). In the C-H region (2850-2990 $cm^{-1}$) the 2979 $cm^{-1}$ is reduced in the silica precipitate, indicating changes in the lysine $CH_2$-group close to the functional amine. A new shoulder at 2850 $cm^{-1}$ could result from a symmetric stretch in $CH_2$- groups (Howell *et al.*, 1999). The sharp peak at 1431 $cm^{-1}$ is assigned to deformations of -$NH_3^+$on the lysine functional group (Aliaga *et al.*, 2009). The broadening and shifting of this peak to higher energies (1456 $cm^{-1}$) indicates an interaction between the silica and the lysine functional group. The interaction may occur through hydroxyl and water groups, inhibiting some -$NH_3^+$ bonds deformations and thus broadening the peak. The peak at 838 $cm^{-1}$ may belong to a C-C stretch in the proline ring (Stewart & Fredericks, 1999a). This vibration disappears in the mineral precipitate, suggesting a structural change in prolines possibly embedded within the silica. Vibrations at 924, 726, and 598 $cm^{-1}$ are assigned to groups of -$COO^-$ at the peptide C-terminal (Stewart & Fredericks, 1999b). These groups seem to participate in the structure of the mineral, as their vibrations are missing in the precipitate.

**Fig. S1** Transcript level of RNA recognition motif-containing protein (*RRM*) in relation to ubiquitin-conjugating enzyme (*UbCE*) as internal control gene, showing that the transcript level of the two house-keeping genes do not change significantly in the tested tissues The error bars represent standard deviation (n=3). The differences in transcript levels among the tissues are statistically insignificant.

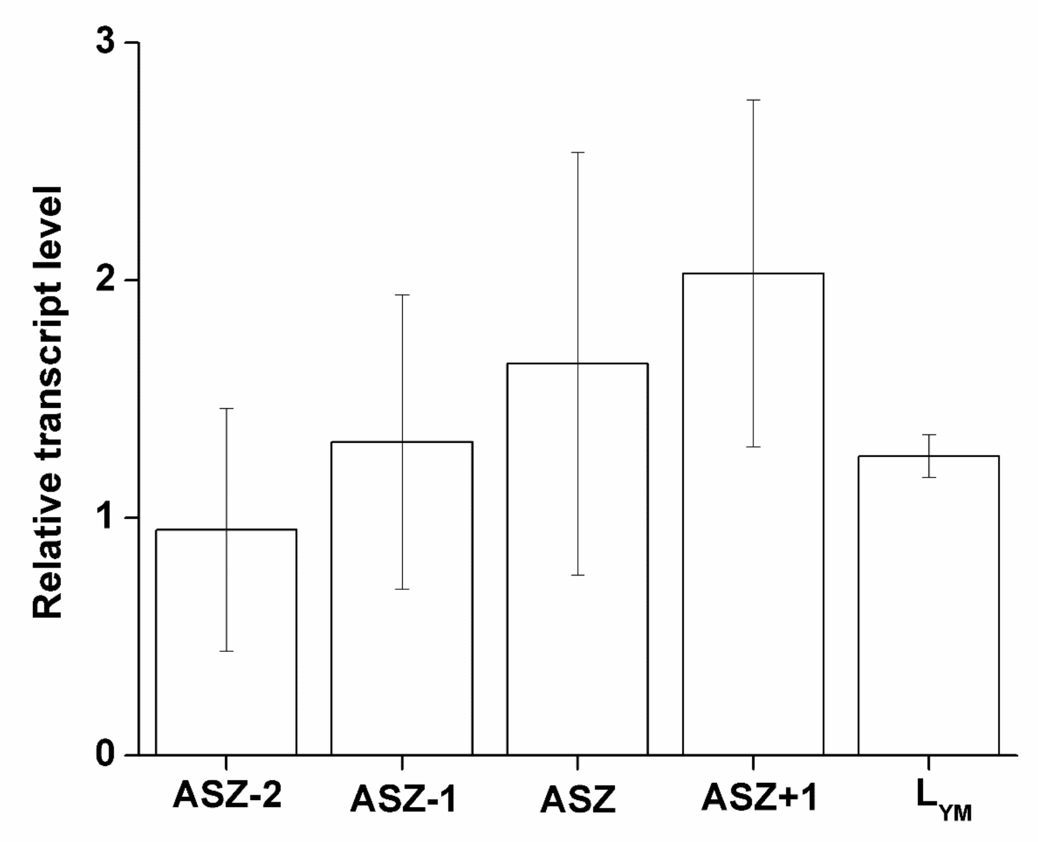

**Fig. S2** Immunolocalization control reactions using pre-immune serum; or lacking either one or both the antibodies. Images from the green channel (fluorophore conjugated with the secondary antibody), red channel (from propidium iodide, which stains the cell wall) and the corresponding white light are merged together. Silica cells are marked with arrows. In any of the control reactions, we did not observe green fluorescence in the tissues. Bar, 50 µm. (a-d) Active silicification zone of wild type sorghum immature, silicifying leaf incubated (a) with pre-immune serum, (b) without anti-Peptide-1 antibody, (c) without secondary antibody, (d) without both the antibodies. (e-h) Leaf pieces from the youngest mature leaf in sorghum, in a zone where all silica cells are already silicified and dead incubated (e) with pre-immune serum, (f) without anti-Peptide-1 antibody, (g) without secondary antibody, (h) without both the antibodies.

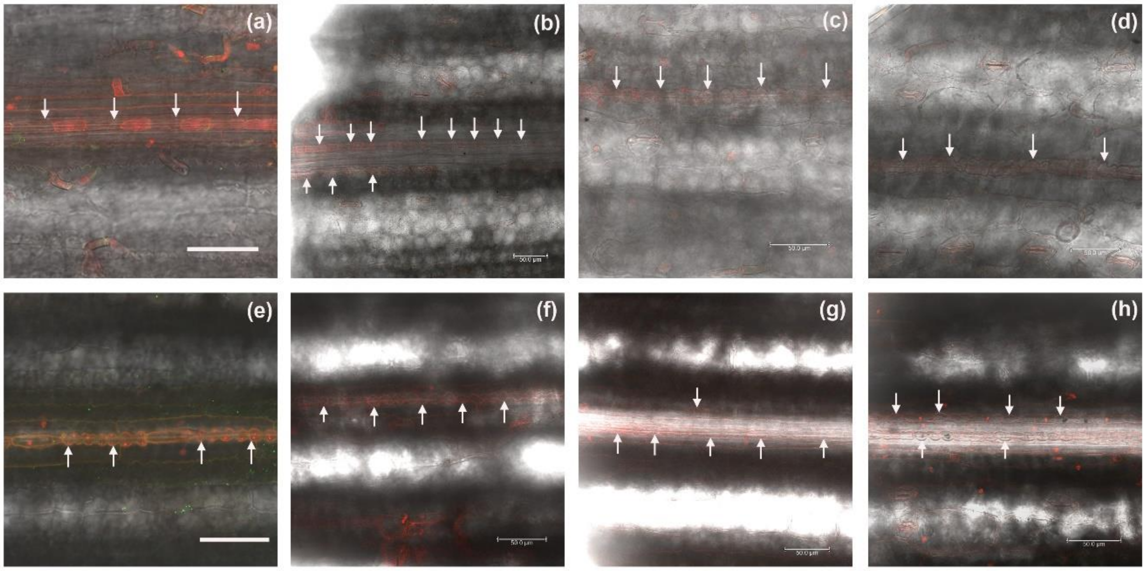

**Fig. S3** Maize dwarf mosaic virus (MDMV) infected sorghum plants showing symptoms. (a) Sorghum plant showing MDMV symptoms (arrow) 3 weeks are after bombardment of MDMV-Slp1 plasmid. (b) Sorghum plant showing MDMV symptoms (arrow) 3 weeks are after bombardment of MDMV-GUS plasmid.

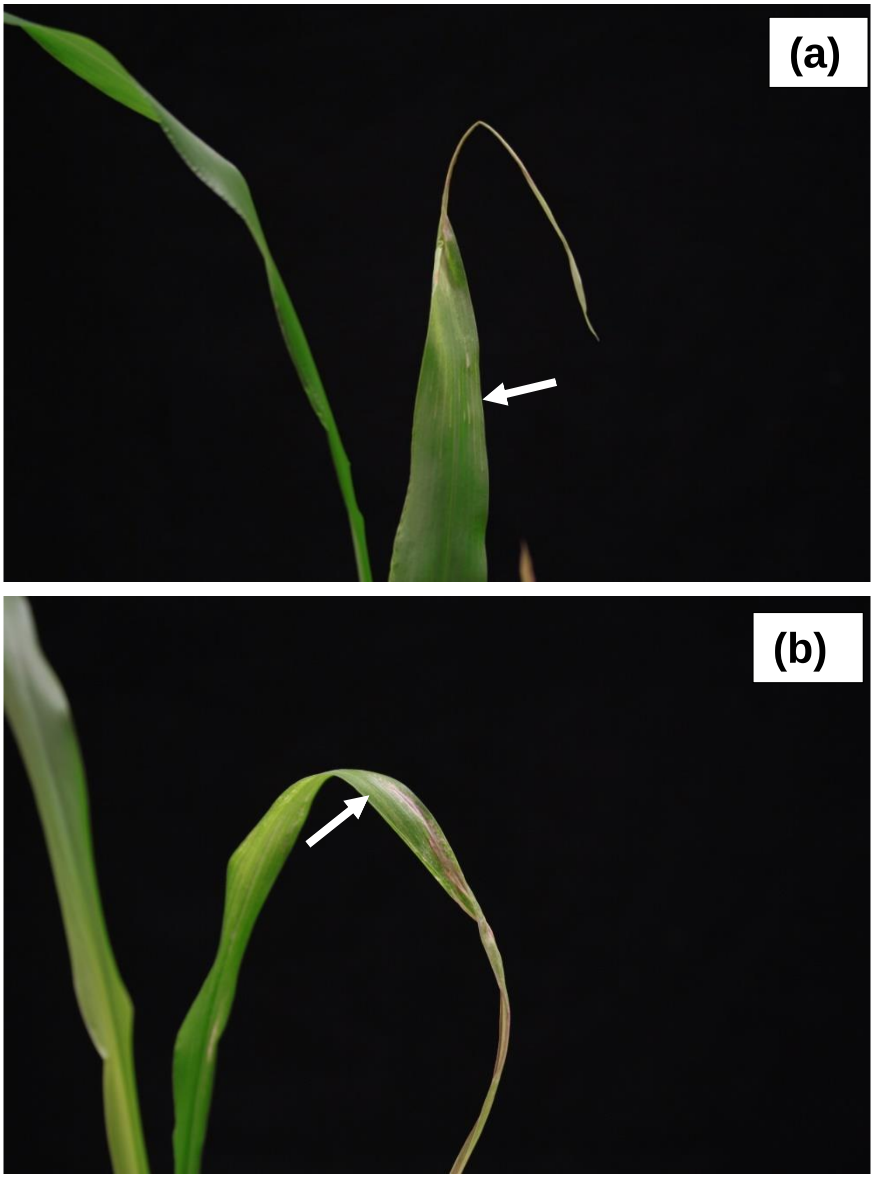

**Table S1** List of primers along with their nucleotide sequence used in the present study. Letters 'F' or 'R' at the end of primer name denote forward and reverse primers, respectively.

| **Primer name** | **Primer nucleotide sequence (5'⟶3')** |
|---|---|
| Slp1-RT-F | TTTCGCCGTGCTGATGGTAA |
| Slp1-RT-R | AAGACCCTTGAATGCCGCTT |
| UbCE-RT-F | CCTTCGGGTCGGTGCTATTT |
| UbCE-RT-R | CGTAAACCTTCAGCAGCCAC |
| RRM-RT-F | CTGAACGAGCGTTGTCGTTG |
| RRM-RT-R | AGCACGCCCTCTAAAGGAAC |
| Slp1-SalI-F | TTTTTTGTCGACATGGCTGCGGTACATGGG |
| Slp1-XbaI-R | TTTTTTTCTAGAGTTTTTGGCCGGAGGATG |
| Slp1-AgeI-F | TTTTTTACCGGTGCTGCGGTACATGGGGGC |
| Slp1-ApaI-R | TTTTTTGGGCCCGTTTTTGGCCGGAGGATG |
| MDMV-Slp1-F | CCGAGGAAGACTTAAAGGCG |
| MDMV-Slp1-R | ATCGTAGGTGTGTGCTCTGT |
| MDMV-Coat-F | AACCCTCGCGACGTTAGAAG |
| MDMV-Coat-R | TGCATAGCATTTCGCCACAC |

**Table S2** List of the sorghum genes identified by BLAST to homologous genes in wheat that show significantly differential transcription upon silicon treatment verses the non-treated plants. Gene id for sorghum Slp1 is shaded in yellow.

| | **Wheat Locus identifier** | **Sorghum Locus identifier** | **Annotations in sorghum as retrieved from Phytozome database** |
|---|---|---|---|
| 1 | Ta#S12989242 | Not found | |
| 2 | Ta#S18011106 | Sb01g048810 | DET1- AND DDB1-ASSOCIATED PROTEIN 1, Det1 complexing ubiquitin ligase |
| 3 | Ta#S12974713 | Sb02g042750 | 60S RIBOSOMAL PROTEIN L18 |
| | | Sb01g035860 | Putative uncharacterized protein |
| | | Sb09g004290 | 60S RIBOSOMAL PROTEIN L18, Ribosomal protein L18e/L15 |
| 4 | Ta#S13208508 | Sb01g005800 | |
| 5 | Ta#S13177960 | Not found | |
| 6 | Ta#S12989467 | Sb03g002980 | |
| 7 | Ta#S13202081 | Sb07g022480 | Family and subfamily not named |
| 8 | Ta#S12995082 | Sb01g041770 | matairesinol biosynthesis, justicidin B biosynthesis |
| 9 | Ta#S12995144 | Not found | |
| 10 | Ta#S13138044 | Not found | |
| 11 | Ta#S13078327 | Sb06g014740 | F28N24.16 PROTEIN, Pollen proteins Ole e I like |
| | | Sb02g012930 | Pollen proteins Ole e I like, Family and subfamily not named |
| | | Sb03g001020 | Pollen proteins Ole e I like, Family and subfamily not named |
| | | Sb09g007260 | Pollen proteins Ole e I like, Family and subfamily not named |
| 12 | Ta#S13156488 | Not found | |
| 13 | Ta#S13139348 | Not found | |
| 14 | Ta#S13156519 | Not found | |
| 15 | Ta#S13234430 | Not found | |
| 16 | Ta#S13166129 | Sb02g037260 | TUBULIN ALPHA-1 CHAIN-RELATED, Tubulin/FtsZ family, GTPase domain |
| | | Sb01g006650 | TUBULIN ALPHA-2 CHAIN-RELATED, Tubulin/FtsZ family, GTPase domain |
| | | Sb01g009560 | TUBULIN ALPHA-2 CHAIN-RELATED, Tubulin/FtsZ family, GTPase domain |
| | | Sb01g009570 | TUBULIN ALPHA-2 CHAIN-RELATED, Tubulin/FtsZ family, GTPase domain |
| | | Sb01g042650 | TUBULIN, Tubulin/FtsZ family, GTPase domain |
| 17 | Ta#S13107961 | Not found | |
| 18 | Ta#S13063289 | Not found | |
| 19 | Ta#S18008495 | Sb01g025970 | Pollen proteins Ole e I like, Family and subfamily not named |
| | | Sb01g026170 | Pollen proteins Ole e I like, Family and subfamily not named |
| | | Sb01g026180 | Pollen proteins Ole e I like, Family and subfamily not named |
| 20 | Ta#S13158129 | Not found | |
| 21 | Ta#S12955535 | Not found | |
| 22 | Ta#S13123378 | Not found | |
| 23 | Ta#S13191618 | Not found | |
| 24 | Ta#S13231500 | Not found | |

**Video/Movie S1** Confocal microscopy video clip showing Slp1 localization in the cytoplasmic space and near the cell boundary of silica cells.

**Video/Movie S2** Confocal microscopy video clip showing the vesicles packed Slp1 fusing to the cell membrane. Slp1 fused to GFP was transiently overexpressed in tobacco leaves. Image stacks were segmented using the program Imaris. Identified vesicles were pseudocoloured red, that can be seen fusing to the cell membrane. GFP fluorescence in the cell margin is also visible in the video.
